# Proximate causes of infertility and embryo mortality in captive zebra finches

**DOI:** 10.1101/847103

**Authors:** Yifan Pei, Wolfgang Forstmeier, Daiping Wang, Katrin Martin, Joanna Rutkowska, Bart Kempenaers

## Abstract

Some species show high rates of reproductive failure, which is puzzling because natural selection works against such failure in every generation. Hatching failure is common in both captive and wild zebra finches (*Taeniopygia guttata*), yet little is known about its proximate causes. Here we analyze data on reproductive performance (fate of >23,000 eggs) based on up to 14 years of breeding of four captive zebra finch populations. We find that virtually all aspects of reproductive performance are negatively affected by inbreeding (mean r = -0.117), by an early-starting, age-related decline (mean r = -0.132), and by poor early-life nutrition (mean r = - 0.058). However, these effects together explain only about 3% of the variance in infertility, offspring mortality, fecundity and fitness. In contrast, individual repeatability of different fitness components varied between 15% and 50%. As expected, we found relatively low heritability in fitness components (median: 7% of phenotypic, and 29% of individually repeatable variation). Yet, some of the heritable variation in fitness appears to be maintained by antagonistic pleiotropy (negative genetic correlations) between male fitness traits and female and offspring fitness traits. The large amount of unexplained variation suggests a potentially important role of local dominance and epistasis, including the possibility of segregating genetic incompatibilities.

## Introduction

Reproductive performance, including offspring survival, is subject to strong directional selection in every generation. Such strong selection works not only on individuals that live in their natural habitat, but also on those that live in captivity, unless artificial selection counters it. Thus, it is puzzling that some populations (or species) have substantial difficulties with successful reproduction, shown as high rates of infertility or embryo mortality. Prominent examples of frequent reproductive failure include humans (De Braekeleer and Dao 1991; Sierra and Stephenson 2006; Miyamoto et al. 2012), and other animals both in natural environments (Lyon 1986; Grossen et al. 2012) and in captive conditions (Ayalon 1978; Bunin et al. 2008; Gwaza et al. 2016; Griffith et al. 2017). Given that directional selection constantly removes genetic variants that lead to poor performance, one might suspect that reproductive failure typically results from inbreeding (Briskie and Mackintosh 2004), because selection against recessive deleterious mutations is inefficient, or from environmental factors (Jurewicz et al. 2009), such as pollutants (Jackson et al. 2011). However, the range of possible explanations is much wider.

Reproductive failure and individual survival are complex traits and hence may be influenced by multiple genetic components that can be evolutionary stable. For instance, reproductive failure and mortality may be caused by selfish genetic elements that are self-promoting at the cost of organismal fitness (Sandler et al. 1959; Lyon 1986; Safronova and Chubykin 2013; Lindholm et al. 2016). Additive genetic variants can also be preserved under intra-locus sexual antagonism, where genes that are beneficial to one sex impose detrimental effects on the other (Foerster et al. 2007; Van Doorn 2009; Innocenti and Morrow 2010). Furthermore, there might be evolutionary trade-offs between traits, such that individuals that invest more in reproduction might show lower survival rates (Stearns 1989; Schluter et al. 1991). A few recent genetic and genomic studies detected genetic variants (e.g. specific genes) involved in dominance effects or rare variants that show main effects on reproductive traits (e.g. Christians et al. 2000; Safronova and Chubykin 2013; Kim et al. 2017; Knief et al. 2017). As an extreme example, a balanced lethal system was identified in crested newts *Triturus cristatus*, where all embryos that are homozygous for chromosome 1 (about 50% of all embryos) die during development (Sims et al. 1984; Grossen et al. 2012).

Despite the development of new genomic tools, it remains difficult to identify and examine the genetic components that show antagonistic effects, or involve more than one locus, i.e. intra- and inter-locus genetic incompatibilities (Dobzhansky 1936; Fishman and Willis 2006; Johnson 2008; Eroukhmanoff et al. 2016). This difficulty is likely due to the complexity of interactions between multiple loci and between the genotype and the environment (Carrell and Aston 2011; Krausz and Riera-Escamilla 2018). If animals in captivity show high rates of reproductive failure because they are not adapted to a given artificial environment, selection can act on the standing genetic variance. This would result in a transient phase where fitness is heritable until the population is better able to cope with the new environment (e.g. due to behavioural and physiological adaptations to captivity). In general, the genetic basis of reproductive failure and variation in survival remains largely unclear in most species.

The zebra finch is a good model species to study how survival and reproductive performance of the two sexes are correlated at the additive genetic level. The zebra finch is a short-lived songbird that easily breeds in captivity (Zann 1996), and its reproductive performance varies extensively among individuals under controlled breeding conditions in both domesticated and recently wild-derived populations (Griffith et al. 2017; Wang et al. 2017). In the wild, the rate of hatching failure (infertile eggs and dead embryos) was estimated to be >15% (table 1). This excludes clutches that failed completely, because nest desertion cannot be ruled out as the reason of failure. In lab stocks, the average proportion of eggs remaining apparently unfertilized ranged from 17% in aviary breeding to 30-35% in cage breeding (table 1), while average embryo mortality rates varied between 24% and 75% (table 1). Average nestling mortality rates were also high (table 1). Although some of the variation has been explained by specific treatment effects (e.g. inbreeding, force-pairing, maternal stress; Hemmings et al. 2012; Ihle et al. 2015; Khan et al. 2016), the high baseline levels of infertility, embryo and nestling mortality remain largely unexplained.

**Table 1:** Summary of rates of hatching failure, infertility, and embryo and offspring mortality reported in the zebra finch literature

| Population | Sample description | Hatching<br>failure (%) | Infertility<br>(%) | Embryo<br>mortality (%) | Nestling<br>mortality (%) | Reference |
| --- | --- | --- | --- | --- | --- | --- |
| Wild | 1,156 eggs, clutches that<br>produced no nestlings were<br>removed | >17 | - | - | - | Zann 1996 |
| Wild | 872 eggs, clutches that<br>produced no nestlings were<br>removed | 16 | - | - | 9 | Griffith et al. 2008 |
| La Trobe University,<br>Australia, domesticated | 31 untreated and<br>25 CORT treated pairs, clutches<br>that produced no nestlings and<br>all first eggs were removed. | Untreated: 24;<br>treated: 45 | Untreated: 7;<br>treated: 15 | Untreated: 10;<br>treated: 29 | - | Khan et al. 2016 |
| Max Planck Institute for<br>Ornithology, Germany,<br>domesticated (from<br>Sheffield, UK) | 11,617 eggs | - | 30 | - | - | Knief et al. 2015b |
| Max Planck Institute for Ornithology, Germany recently wild-derived (from Bielefeld, Germany) | 852 eggs, aviary | - | 17 | 24 | 45 | Ihle et al. 2015 |
| Sheffield University, UK | 161 eggs for infertility; 2,884 eggs for hatching failure and nestling mortality | 52 | 35 | - | 31 | Kim et al. 2017 |
| Sheffield University, UK | 1,524 eggs; 77 unrelated and 20 sib-sib pairs | - | Unrelated: 9;<br>sib-sib: 11 | Unrelated: 59;<br>sib-sib: 75 | Unrelated: 55;<br>sib-sib: 68 | Hemmings et al. 2012 |

Note: For population of La Trobe University, Australia, in treated pairs, females were given a corticosterone (CORT) mix after laying the first egg. The CORT mix was made of 0.5 mg crystalline corticosterone dissolved by 10 µl ethanol then mixed with 990 µl peanut oil (Khan et al. 2016). Hatching failure: proportion of eggs that do not hatch. Infertility: proportion of eggs that show no sign of development. Embryo mortality: proportion of fertilized eggs where the embryo died before hatching. Nestling mortality: proportion of nestlings that died before fledging.

To better understand this variation in reproductive performance and individual survival, we here report on a comprehensive quantitative genetic analysis of lifespan, fecundity, infertility, offspring mortality and other fitness-related traits that cover most phases of reproduction for the two sexes (table 2). We quantified the effects of inbreeding, age and an individual’s early nutritional condition on all measured aspects of reproductive performance and survival.

**Table 2:**
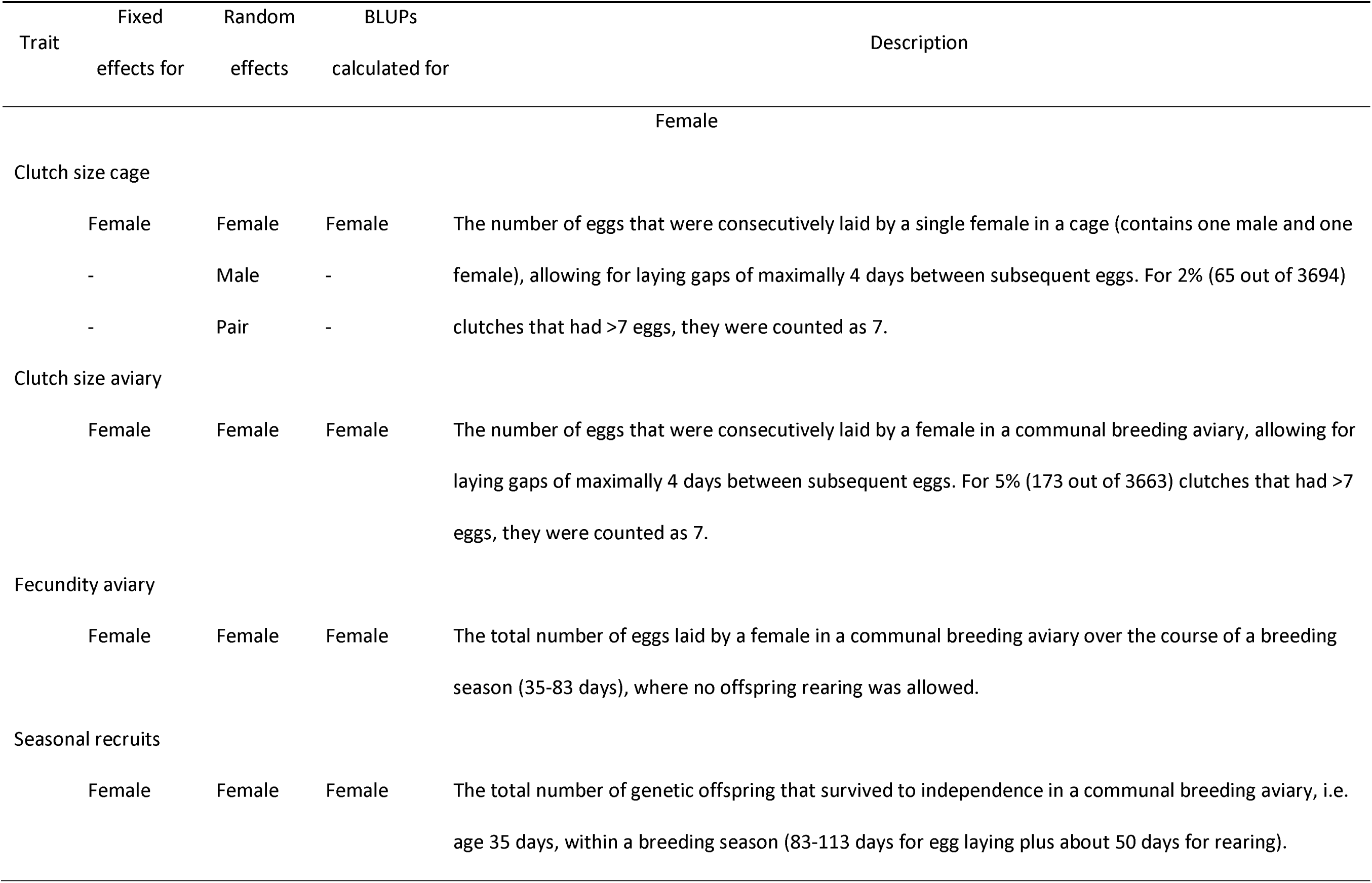
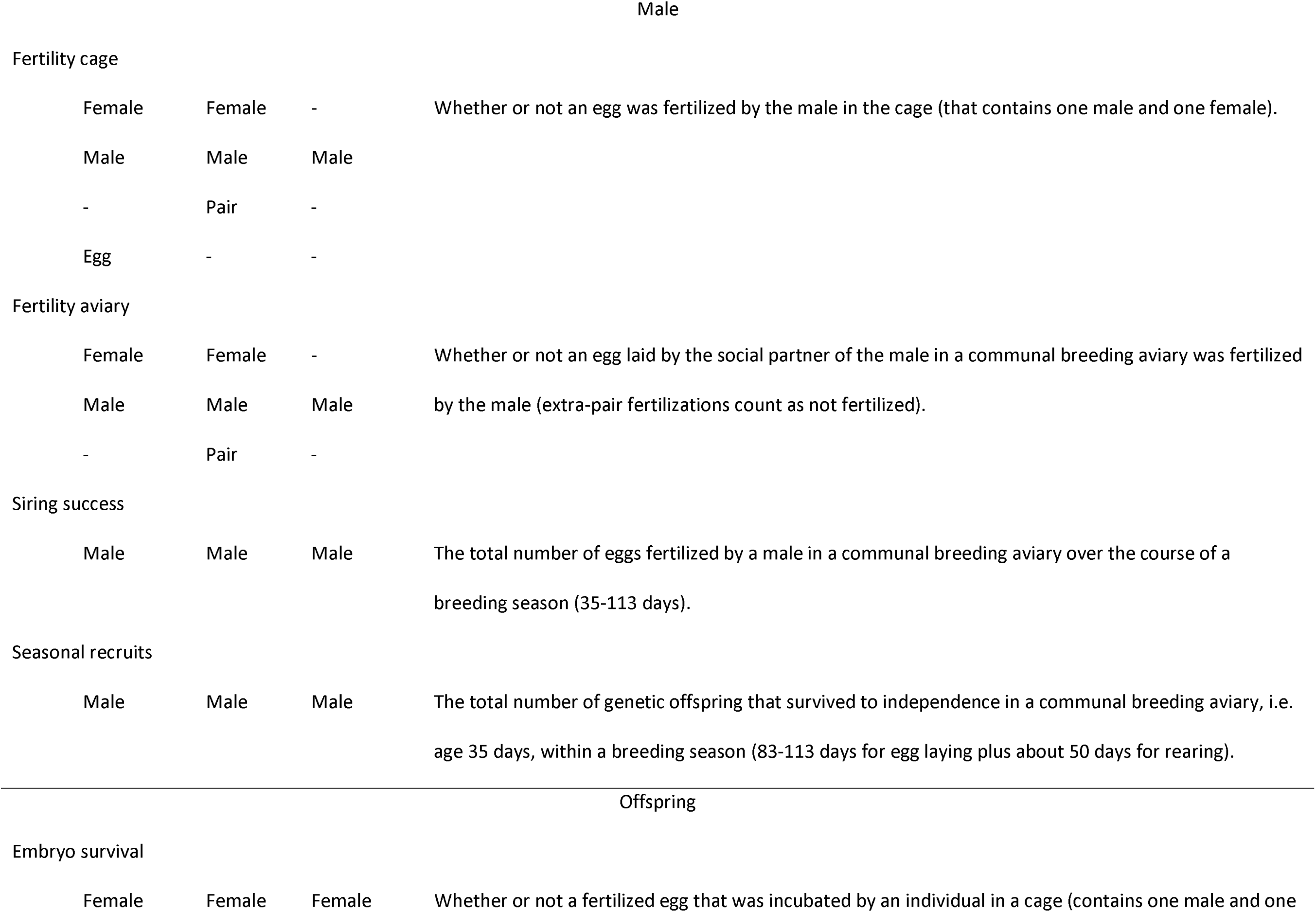
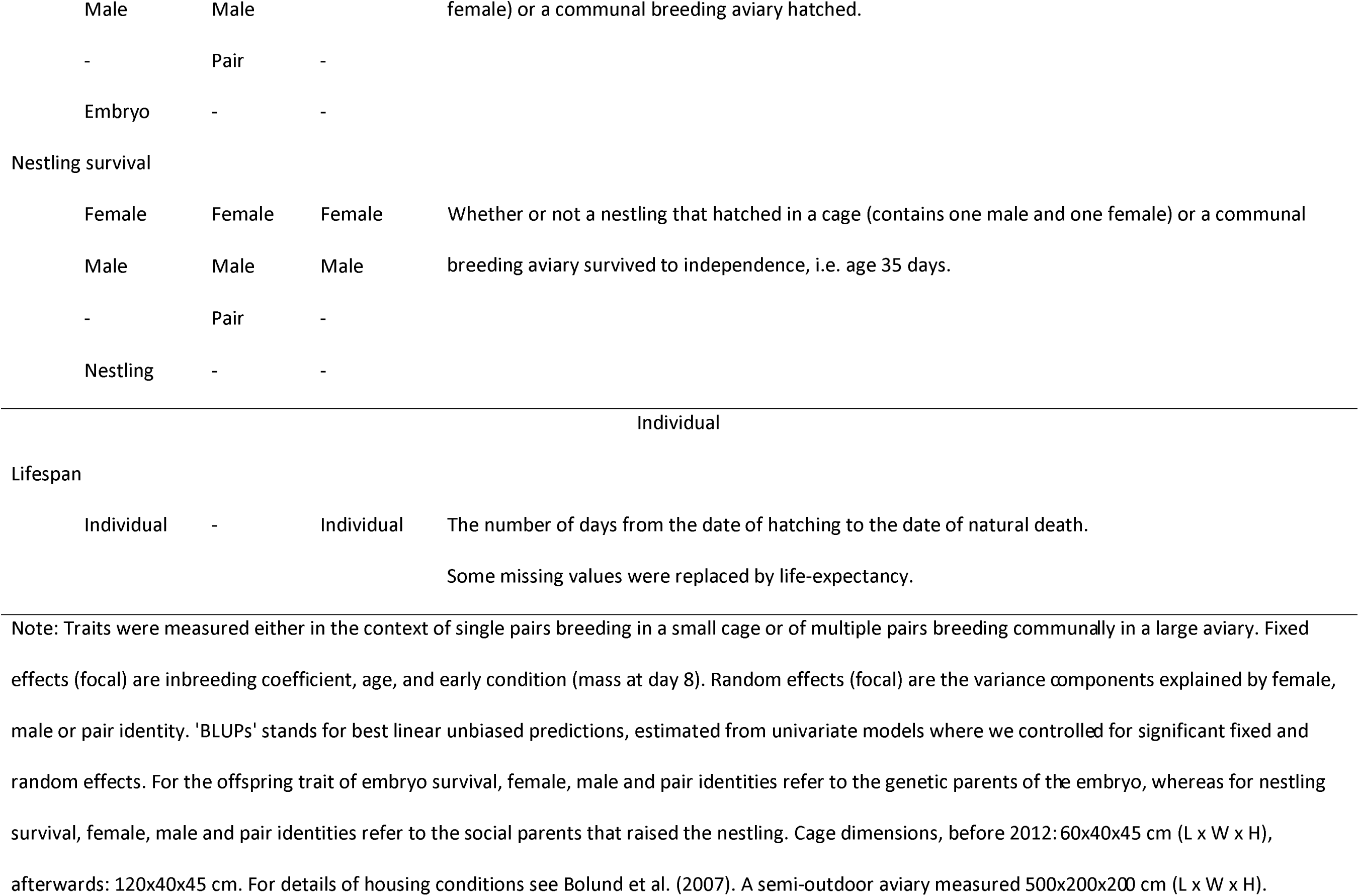
Description of reproductive performance traits in our zebra finch study

Wild zebra finches have a remarkably large effective population size (Balakrishnan and Edwards 2009), where inbreeding is almost completely absent (Knief et al. 2015a). In contrast, in captivity, mating between related individuals is practically inevitable in the long run (Knief et al. 2015*a*). The level of inbreeding typically correlates negatively with individual fitness and various morphological and life-history traits, even though the estimated effect sizes can vary widely (Charlesworth and Charlesworth 1987; Keller and Waller 2002; Bolund et al. 2010a; Forstmeier et al. 2012; Hemmings et al. 2012; Hoffman et al. 2014; Huisman et al. 2016; Michaelides et al. 2016). The importance of inbreeding in predicting reproductive failure remains largely unclear.

Ageing, or senescence, typically leads to a decline in reproductive function at old age, e.g. in birds (Bouwhuis et al. 2009; Lecomte et al. 2010) and humans (Speroff 1994; Shirasuna and Iwata 2017). In zebra finches breeding in cages, male and female fertility declined when individuals became older (Knief et al. 2017). More generally, the relationship between age and reproductive performance is often quadratic, with an initial increase in performance due to gained experience that may mask any early-starting decline caused by deterioration of the body (Harely 1990; Bouwhuis et al. 2009; Lecomte et al. 2010).

The conditions that an individual experienced during early development may also affect fitness later in life. Such permanent environmental effects have been demonstrated using brood size manipulations and they may affect individual behavior and reproductive investment (Gorman and Nager 2004; Tschirren et al. 2009; Rickard et al. 2010; Boersma et al. 2014). In zebra finches, being raised in enlarged broods apparently did not affect later performance (Tschirren et al. 2009). However, a non-experimental measure of individual early-growth condition, namely body mass measured at 8 days of age (which ranges from 2-12 grams), had a significant but small effect on fitness later in life (Bolund et al. 2010*b*).

For this study, we used systematically recorded data on individual body mass at 8 days of age and on reproductive parameters and survival for four captive populations of zebra finches with an error-free pedigree. The aims of this study were (1) to estimate and compare the relative importance of inbreeding, early nutritional condition and age on reproductive performance and lifespan, (2) to estimate the relative importance of individual and pair identity (i.e. repeatability) on reproductive performance, (3) to quantify the heritability of individual reproductive performance and (4) to test if some of the heritable components can be maintained by antagonistic pleiotropy, by analyzing the additive genetic correlations between reproductive performance traits and lifespan across the two sexes.

## Methods

Zebra finches are opportunistic breeders that are abundant throughout most of Australia. Individuals become sexually mature around the age of 90 days and then form pairs for life through mutual mate choice. Breeding pairs cooperatively incubate and raise nestlings until they reach independence around the age of 35 days (Zann 1996). Captive zebra finches live for about 4.5 years on average and maximally for 10 years (Zann 1996, our unpublished data). The studied zebra finches originated from four populations held at the Max Planck institute for Ornithology, Seewiesen, Germany. The population background, rearing conditions and breeding seasons have been detailed elsewhere (see also the online appendix, tables A1 and A2). In brief, we compiled and analyzed up to 14 years of zebra finch reproductive performance data from

1. population ‘Seewiesen’, a domesticated population derived from the University of Sheffield, with a nine-generation long error-free pedigree (population #18 in Forstmeier et al. (2007b));
2. population ‘Krakow’, a domesticated population that was generated by hybridizing between Krakow (#11 in Forstmeier et al. (2007b)) and Seewiesen populations;
3. population ‘Bielefeld’, which was derived from the wild in the late 1980s (#19 in Forstmeier et al. (2007b));
4. population ‘Melbourne’, which was derived from the wild in the early 2000s (see Jerónimo et al. (2018)).

Birds from the two recently wild-derived populations were smaller (ca. 11g) compared to domesticated birds (ca. 15-16g), and more shy, so we only bred them in large semi-outdoor aviaries (rather than in small cages, see table 2 for sizes of cage and aviary).

Between 2004-2017, we bred zebra finches in four settings with various treatments (see tables A1 and A2 for details): (1) cage breeding, (2) cage laying, (3) aviary breeding, and (4) aviary laying. In cages, single pairs were kept and hence partners were assigned. In aviaries, groups of birds were kept together and individuals could freely form pairs. Group size was typically 12, but ranged from 10 to 42, with sex ratio (proportion of males) ranging from 0.4 to 0.6. In a ‘breeding’ setup, pairs were allowed to rear their offspring, whereas in a ‘laying’ setup all eggs were collected for paternity assignment and replaced by plastic eggs that were removed after 7 or 10 days of incubation. The proportion of individuals that participated in more than one breeding season ranged from 0.23-0.84 (mean 0.47).

In this study, we focus on general effects on reproductive performance in zebra finches, not on population-specific effects. Therefore, in all analyses, we only controlled statistically for between-population differences in reproductive performance (main effects only, no interactions).

### Measures of the focal fixed effects: inbreeding, age and early nutrition

We used the pedigree-based inbreeding coefficient ‘F_ped_’, calculated using the R package ‘pedigree’ V1.4 (Coster 2015), as a measure of the degree of inbreeding of an individual (Wright 1922; Knief et al. 2015a). F_ped_ reflects the proportion of an individual’s genome that is expected to be identical by descent (Howrigan et al. 2011; Knief et al. 2015a). For instance, full-sibling mating produces inbred offspring that are expected to have 25% of the genome identical by descent (F_ped_ = 0.25). For practical reasons, all founders were assumed to be unrelated (F_ped_ = 0; Forstmeier et al. 2004). However, their true level of identity by descent is likely about 5% (judging from runs of homozygosity; Knief et al. 2015a).

For all birds, we recorded their exact hatch date. Thus, for models of reproductive performance at the level of eggs, clutches, and breeding rounds (as the unit of analysis), we used the exact age (in days) of the female or the male when an egg was laid, a clutch started, or a breeding round started, respectively. At the start of reproduction, individuals were 69-2909 days old. On the day of hatching, we individually marked all nestlings on the back using water-proof marker pens (randomly using red, blue and green, and pairwise combinations of these colors if >3 nestlings). We checked survival almost daily (daily on weekdays, occasionally during weekends) until offspring became independent (age 35 days). As a measure of early-growth condition, we determined body mass of each nestling to the nearest 0.1 g at 8 days of age (hereafter ‘condition’). Despite the fact that high-quality food was available to all parents *ad libitum*, nestling body mass at this age ranged from about 1.5 g to 12.6 g (mean = 7.1 ±1.7 SD). For 297 out of 6190 nestlings, body mass was measured on day 6, 7 or 9. For those individuals, we estimated their mass on day 8, as follows. We constructed a linear mixed-effect model, with nestling body mass as the dependent variable, with the actual age of the mass measurement as a continuous covariate and with F_ped_ and population (1-4, see above) as fixed effects. We also included the identity of the genetic mother as a random effect. Using the slope of daily mass gain, we estimated mass at day 8 for those 297 individuals by adding or subtracting 0.97 g per day of measuring too early or late. Because the four populations differ in body mass, we normalized (Z-scaled) all measured or estimated values of mass at day 8 within each population before further analysis.

We report effects of inbreeding, age and early condition always with a negative sign, such that negative values of greater magnitude reflect stronger detrimental effects of being inbred, old, or poorly fed. This allows to meta-summarize the results and to directly compare the strength of the focal fixed effects on reproductive performance.

### Measures of lifespan and reproductive performance traits

Table 2 provides an overview of all traits included in this study. To allow direct comparison and easy interpretation of the fixed effects and additive genetic correlations, we scored all traits such that higher, positive values reflect better reproductive performance.

Lifespan was analyzed in the following subset of birds: 5 generations of birds from the Seewiesen population (referred to as generations P, F1-F3, and S3, N = 1855 individuals) and 4 generations of birds from the Bielefeld population (F1-F4, N = 1067 individuals). Among those birds, we used the 4 most complete generations P and F1-F3 Seewiesen for which we recorded the exact lifespan for all (N = 1175 individuals) as a pool to impute missing lifespans. For 219 S3 Seewiesen birds and for 663 Bielefeld birds, no date of natural death was available (e.g. because individuals were still alive or because their fate was unknown). For these individuals, we used imputed life expectancy in all analyses, defined as the average lifespan of individuals from the same pool that lived longer than the focal bird when last observed alive.

In aviaries, we identified social pairs by behavior (clumping, allopreening, and visiting a nest together). All parentage assignments were based on conventional microsatellite genotyping, following Forstmeier et al. (2007a). We assigned every fertilized egg to its genetic mother (N = 11704 eggs). When the egg appeared infertile (no visible embryo; Birkhead et al. 2008), we assigned it to the social female that was attending the clutch (N = 3630 cases). In 36 cases where two females used the same nest to lay eggs, we assigned the unfertilized eggs to the female that laid the most similar eggs (in size and shape), based on eggs that were certainly laid by a given female (e.g. fertilized eggs and eggs in other clutches laid by that female). In cases where birds were not allowed to rear offspring, we quantified female fecundity as the total number of eggs laid by the focal female during the breeding period (see table A1 and A2).

In breeding experiments, we opened all unhatched eggs to check for visible signs of embryo development and classified them as either infertile or ‘embryo mortality’. In experiments in which all eggs were incubated artificially for a few days to collect DNA from embryos, we classified eggs as infertile or not, but discarded information on embryo viability. Visual inspection of opened eggs has the disadvantage that early embryo mortality may get misclassified as infertility if it occurred before any visible signs of development. Misclassification cannot be avoided entirely, even with more time-consuming examination of eggs, which would be challenging to do for thousands of eggs (Bellairs and Osmond 2005; Birkhead et al. 2008; Murray et al. 2013). However, most cases of apparent infertility coincided with the absence of sperm on the perivitelline layer of the egg (fig. A1, see also Birkhead and Fletcher (1998)). Thus, we expect only a small fraction of misclassification.

In cages, we measured male fertility as a binary trait, i.e. whether an egg was fertilized or not. In 12 cases, one to five eggs (median: 1 egg) were fertilized by the previous partner of the female and those were counted as infertile eggs of the focal male. In aviaries, we assessed fertility by whether an egg that was laid by a male’s social partner was sired by him or not. Thus, in aviary conditions, fertility also reflects a male’s ability to defend his paternity against extra-pair males. We also quantified male siring success as the total number of fertilized eggs sired by a focal male. This includes males that remained unpaired (without a social female).

For each fertilized egg that was incubated by the social parents, we recorded whether it hatched or not (binomial trait for the genetic parents). For each hatched egg that was reared, we recorded whether the nestling survived to independence (day 35; binomial trait for the social parents). We quantified the number of seasonal recruits as the number of genetic offspring that survived to independence within a given breeding season. Number of seasonal recruits was square-root transformed to approach normality.

### Statistical models

All mixed-effect models were run in R, using the R packages ‘lme4’ V 1.1-18-1 (Bates et al. 2018). All animal models were run using VCE6 (Neumaier and Groeneveld 1998), because (a) it allows running a 12-trait multivariate animal model that consists of 2346 individuals with at least one trait value per individual and (b) it has a reasonable running time. To check the consistency of model outputs, we repeated all animal models in the R packages ‘pedigreemm’ V 0.3-3 (Vazquez et al. 2010; univariate animal models only) and ‘MCMCglmm’ (Hadfield 2015; univariate and bivariate animal models). All model details are listed with the supporting data and R scripts at https://osf.io/tgsz8/. Model outputs of all methods are given in the online appendix. The heritability and additive genetic correlation estimates were highly correlated between methods (r>0.65, P<0.002). We report the VCE6 estimates, unless otherwise stated. Figure A2 shows the exact range of each focal fixed effect and each performance trait value. Here, we Z-transformed all covariates and response variables across populations to allow direct comparison of the effect sizes for inbreeding, age and condition across all models. The 95% CIs of fixed effects from mixed-effect models were calculated using the function ‘glht’ from the R package ‘multcomp’ V1.4-10 while controlling for multiple testing (Hothorn et al. 2008).

Data analysis involved four consecutive steps (fig. 1):

**Figure 1.**
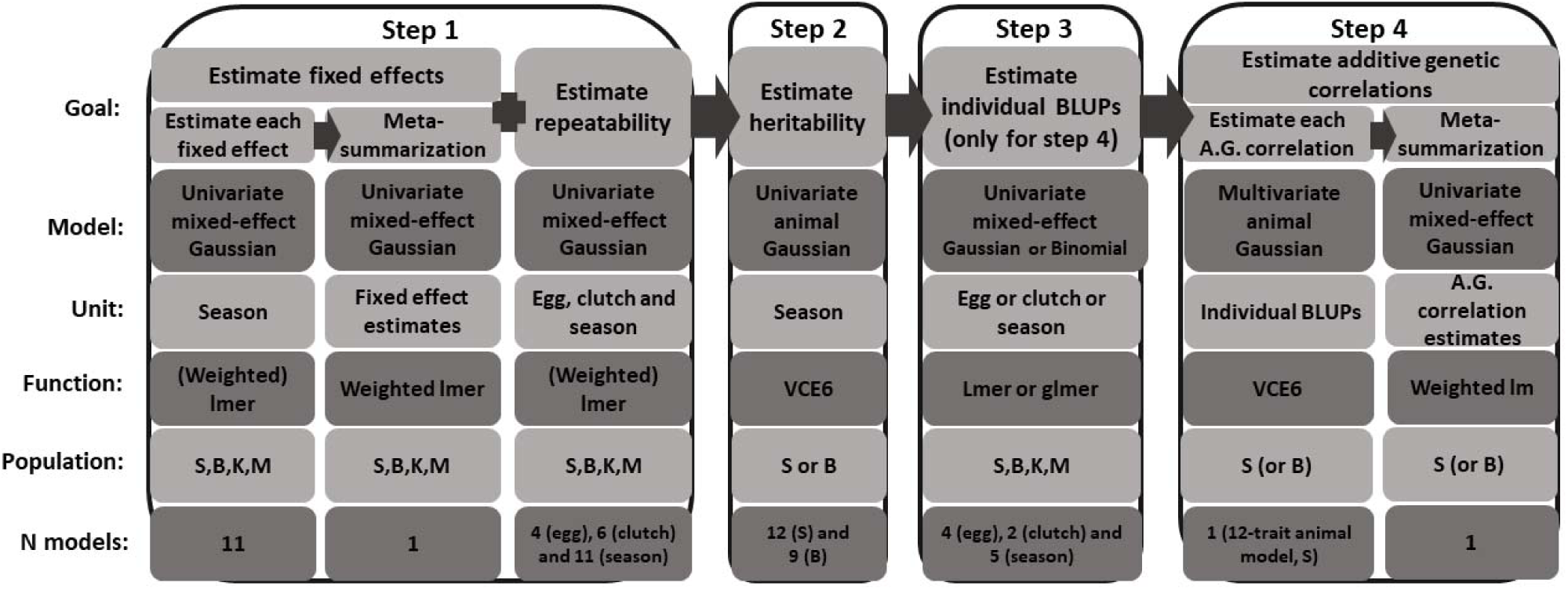
Steps of data analysis from univariate mixed models to multivariate animal models. Shown are the goals of the analysis, the model properties, the unit of analysis (i.e. whether rows in the data represent single eggs, clutches, individuals in a breeding season, single fixed effect estimates, or individuals overall), the software functions used for analysis (for models on aggregated levels, ‘weight’ stands for the number of eggs or clutches used for each aggregation, whereas in meta-summarization models, weight stands for the multiplicative inverse of the standard error of each estimate) and the population abbreviations for data used for the analysis. ‘S’, ‘B’, ‘K’, and ‘M’ stand for ‘Seewiesen’, ‘Bielefeld’, ‘Krakow’, and ‘Melbourne’, respectively. Number of models conducted within each step with their specific details (e.g. unit, population or the model type) used for analysis. ‘A.G.’ stands for ‘additive genetic’.

#### Step 1: Estimation of fixed effects and variance decomposition

The goal of Step 1 was to estimate (a) all fixed effects on reproductive performance and (b) individual repeatability of performance traits. All fixed and random effects of models used in Step 1 are listed in tables A3-A4. In brief, we first fitted all models with a Gaussian error distribution to compare and meta-summarize the estimated effect sizes of the fixed effects and to estimate the variance components for the random effects. We used all observations with information on the three fixed effects (age, F_ped_, and early condition of the male, female and the individual egg if applicable), and included population (fixed effect) and female, male, and pair identity (random effects). We analyzed traits that were measured at either egg, clutch, or season level. As applicable, we fitted as fixed effects the laying sequence of eggs within a clutch, the order of hatching of offspring within a brood, the order of the clutches that were laid by a female over the course of a season, the sex ratio in the aviary, and the duration of the season (table A1). For models of embryo survival, we also controlled for whether or not the eggs were incubated in a nest that still contained offspring from a previous brood (7% of embryos). For models of nestling survival, we added as fixed effect pair type (pair formed through mate choice or through force-pairing; Ihle et al. 2015). For models of egg-based fertility, embryo and nestling survival, we also tested the effect of egg volume on egg fate (we calculated volume as 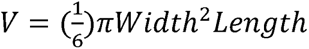 where egg length and width had been measured to the nearest 0.1 mm). For this analysis, we fitted the mean egg volume of each female and the centered egg volumes (centered within individual females) to distinguish between the effects of between- and within-female variation in egg size (van de Pol and Wright 2009). We estimated the variance components for male, female, and pair identity, and further controlled for clutch identity and identity of the setup (see appendix tables A1-A2), as applicable, by adding them as random effects. Lifespan had no repeated measurement, therefore we only included individual identity as a dummy random effect for practical reasons when running the model and extracting estimates in R. For this ‘lm’ model, the correlation between the residuals and the dummy random effect equals one, and the fixed effect estimates were unaffected by the dummy variable. Table 2 shows for which group of individuals, i.e. female, male or the offspring itself, we tested which focal fixed and random effects.

To allow direct comparison of the magnitude of fixed effects at the same level of measurement, we also aggregated data within clutches (e.g. proportion of infertile eggs within a clutch) and within individuals over the course of a season. Models on aggregated data were weighted by the number of eggs within a clutch or by the number of eggs or clutches for an individual within a season (fig. 1). As expected, the proportion of variance explained by male, female and pair identity increased from the egg level to the season level (see Results). However, the relative proportions explained by female, male, and pair identity did not change notably. Therefore, we focus on the analyses of fixed effect estimates at the breeding season level.

To compare the overall effect sizes between the focal fixed effects, we meta-summarized the estimated effect sizes for inbreeding, age and condition using the weighted ‘lmer’ function from the R package ‘lme4’. The uncertainty of each estimate was accounted for by using the multiplicative inverse of the standard error 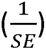 of the response variable as ‘weight’. In this meta-model, we used effect size estimates from models that had been aggregated at the season level as the dependent variable. Note that effects of inbreeding of the egg on fertility in cage-breeding and nestling survival were taken from egg-based models, because they cannot be aggregated by clutch or season. Additionally, we tested whether effect sizes differed between males, females and offspring (fixed effect with three levels) or among traits (random effect with 11 levels; as listed in table 2).

Additionally, we tested for early-starting ageing effects by selecting reproductive performance data for males and females that were <2 years old when reproducing. We then meta-summarized the mean age effect estimates using the R function ‘lm’, weighted by the multiplicative inverse of the standard error.

We calculated the amount of variance explained by each fixed effect (Nakagawa and Schielzeth 2010) as the sum-of-squares of the fixed effect divided by the number of observations (N-1) (Henderson 1953). In weighted models, we divided the variance components of the fixed effects and the residual by the mean weight value (Bates et al. 2018).

#### Step 2: Estimation of heritability of fitness-related traits

The goal of Step 2 was to estimate the heritability of reproductive performance traits using univariate Gaussian animal models. Because quantitative genetic models require large amounts of data, we restrict our analyses to the populations Seewiesen and Bielefeld. Note that the pedigrees of our four captive populations are not connected, so it was not useful to analyze them jointly.

We kept the general model structure from Step 1, but excluded the fixed effects of egg volume on male fertility, embryo and offspring survival (to avoid removing biological variation that is potentially heritable and hence of interest; note that the effect sizes of egg volume are small, see Results). For the embryo survival model, we excluded the non-significant fixed effects of male age, inbreeding, and condition. For the model on male fertility from cage-breeding, we excluded the non-significant effect of the level of inbreeding of the egg itself. To most effectively use the available information on reproductive performance, we included individuals with missing values for condition (N = 231 founder individuals and N = 23 individuals of the F2 generation; i.e. 7% of Seewiesen birds). These missing values were replaced by the population mean. Individual identity was fitted twice, once linked to the individual correlation matrix (pedigree) to estimate the amount of variance from additive genetic effects (V_A_) and once to estimate the remaining amount of variance from permanent environmental effects (V_PE_) (Kruuk and Hadfield 2007). Animal models on nestling mortality were run twice, once for the mother and once for the father. We calculated heritability based on the total phenotypic variance, V_Ph_, as h^2^ = (V_A_/V_ph_), and we also quantified V_A_ relative to the individual repeatability as (V_A_/(V_A_+V_PE_)).

We compared the estimates of heritability (and V_A_ relative to the individual repeatability), between the domesticated population ‘Seewiesen’ and the recently wild-derived population ‘Bielefeld’ using the R function ‘lmer’. We used the multiplicative inverse of the standard error as ‘weight’ to control for variation in uncertainty of each estimate. We used the estimates of heritability as the response variable, and fitted population as fixed effect (two levels) and trait as a random effect (9 levels, only including traits that were measured in both populations).

#### Step 3: Calculation of mean individual fitness-related traits values using BLUPs

The only goal of Step 3 was to extract individual estimates of reproductive performance needed for Step 4. We kept the model structure from Step 1, except that we used a binomial error structure for binary traits, i.e. male fertility in cages and aviaries, embryo and nestling survival. Missing values for condition (mostly founders of each population, 6% of all birds of the four populations) were replaced with population means as in Step 2. For the embryo survival model, we again excluded the non-significant effects of male inbreeding, age, and condition. We also excluded (a) effects of egg volume from all egg-based models and (b) the effect of the level of inbreeding of the egg itself from the model of male fertility measured in cages (see Step 2).

We extracted the best linear unbiased predictions (BLUPs) for female or male identity (as applicable) as the estimated life-history trait value of that individual (table 2) for Step 4.

#### Step 4: Estimation of additive genetic correlations

The goal of Step 4 was to estimate additive genetic correlations between different performance traits using multivariate animal models.

Before fitting a 12-trait animal model that estimates for each matrix (genetic and residual) all 12 variances and 66 covariances simultaneously, we aggregated the raw data to one phenotypic value per individual for each trait. This was necessary because we are not aware of software that can handle the full complexity of the underlying raw data (involving more than 26 different fixed effects). Because simple averages of multiple measures can result in outliers when sample size is small, we used the phenotypic BLUPs described above. BLUPs do not produce outliers and account for all considered fixed and random effects (Robinson 1991; Houslay and Wilson 2017). Breeding values (genetic BLUPs) suffer from non-independence, because the phenotype of one individual influences the breeding values of all its relatives (Hadfield et al. 2010). Note that this is not the case for the phenotypic BLUPs we use here. However, the uncertainty that is inherent to each BLUP is not taken into account, which may lead to underestimation of standard errors (Houslay and Wilson 2017). To check the robustness of our results, we compared our estimates with those obtained (a) using a smaller dataset from another population (‘Bielefeld’) with the same method and (b) using bivariate animal models in ‘MCMCglmm’ V 2.26 (Hadfield 2015) (population ‘Seewiesen’). The latter approach is presumably less powerful than a full 12-trait animal model.

For each of the 12 traits, we fitted an intercept, and the pedigree as the only random effect to separate additive genetic from residual variance. We ran these models for the largest and most comprehensive dataset (population Seewiesen; N = 2346 individuals with at least one trait value, BLUPs for 12 traits, 66 covariances) and for the more limited dataset (population Bielefeld; N = 1134 individuals, BLUPs for 9 traits, 36 covariances; see Results).

We used the weighted ‘lm’ function in the R package ‘stats’ to summarize the estimated additive genetic correlations within and between the major categories of traits, i.e. female, male, offspring traits, and lifespan for each population separately (table 2). We fitted the estimates of additive genetic correlations (for each pair of traits, weighted by the multiplicative inverse of the standard error of each estimate) as the dependent variable with trait-class combination as a predictor with seven levels. We removed the intercept to estimate the mean additive genetic correlation for each pairwise combination of classes. We then computed the eigenvectors of the additive genetic variance-covariance matrix of traits, using the R function ‘eigen’, and visualized the orientation of the traits in the additive genetic variation space defined by the principle components PC1 and PC2.

## Results

### Effects of laying and hatching order, clutch order and egg volume on egg and embryo fate

The fate of an egg and its embryo depended on the order of laying within a clutch, the order of hatching within a brood, and the order of consecutive clutches within a breeding season (fig. A3; table A3, models at the ‘Egg’ level). First-laid eggs in a clutch were significantly more likely to be infertile or to contain a dead embryo. Fertility and embryo viability were the highest for the 3^rd^ egg (fig. A3). Male fertility significantly increased over the first three clutches and stayed high afterwards. In contrast, clutch order did not affect the probability of embryo and nestling survival.

The average effect of egg volume on measures of egg fate was small (mean r = 0.040±0.016 SE, fig. A4). Effects of egg volume were largest for nestling survival after hatching, and smallest for embryo survival (table A3, fig. A4). Despite large sample size (N = 9,785 eggs), embryo survival was not significantly influenced by egg volume (between-female variation: r = 0.015±0.017 SE, P = 0.37; within-female variation: r = 0.018±0.010 SE, P = 0.08; table A3). Additionally, embryos in clutches that were incubated in the presence of nestlings from previous breeding attempts were more likely to die before hatching (b = 0.192±0.048 SE, P < 0.0001; table A3). Overall, the total amount of variance explained by laying and hatching order, clutch order and egg volume on egg fate was less than 5% (table A4).

### Effects of inbreeding, age, and early condition

Individuals performed worse in virtually all studied reproductive traits when they were more inbred, as they became older and when they weighed less at 8 days of age (figs. 2, A2 and A5, table A3). Interestingly, reproductive performance did not show an initial increase at young age (meta-summarized effect size of age among birds younger than 2 years: r = -0.013 ± 0.011 SE, figs. 2C, F, 3 and A2). Inbred eggs were equally likely to be infertile than outbred eggs, while inbred embryos and offspring were more likely to die (fig. 3C). Together, this suggests that most infertile eggs were not cases of undetected early embryo mortality. Individuals lived shorter when they were inbred and when they had low weight at day 8 (fig. 3, table A3). However, the fixed effects of inbreeding, age and condition together explained on average only 2% of the variance across all traits (fig. 4 and table A5).

**Figure 2.**
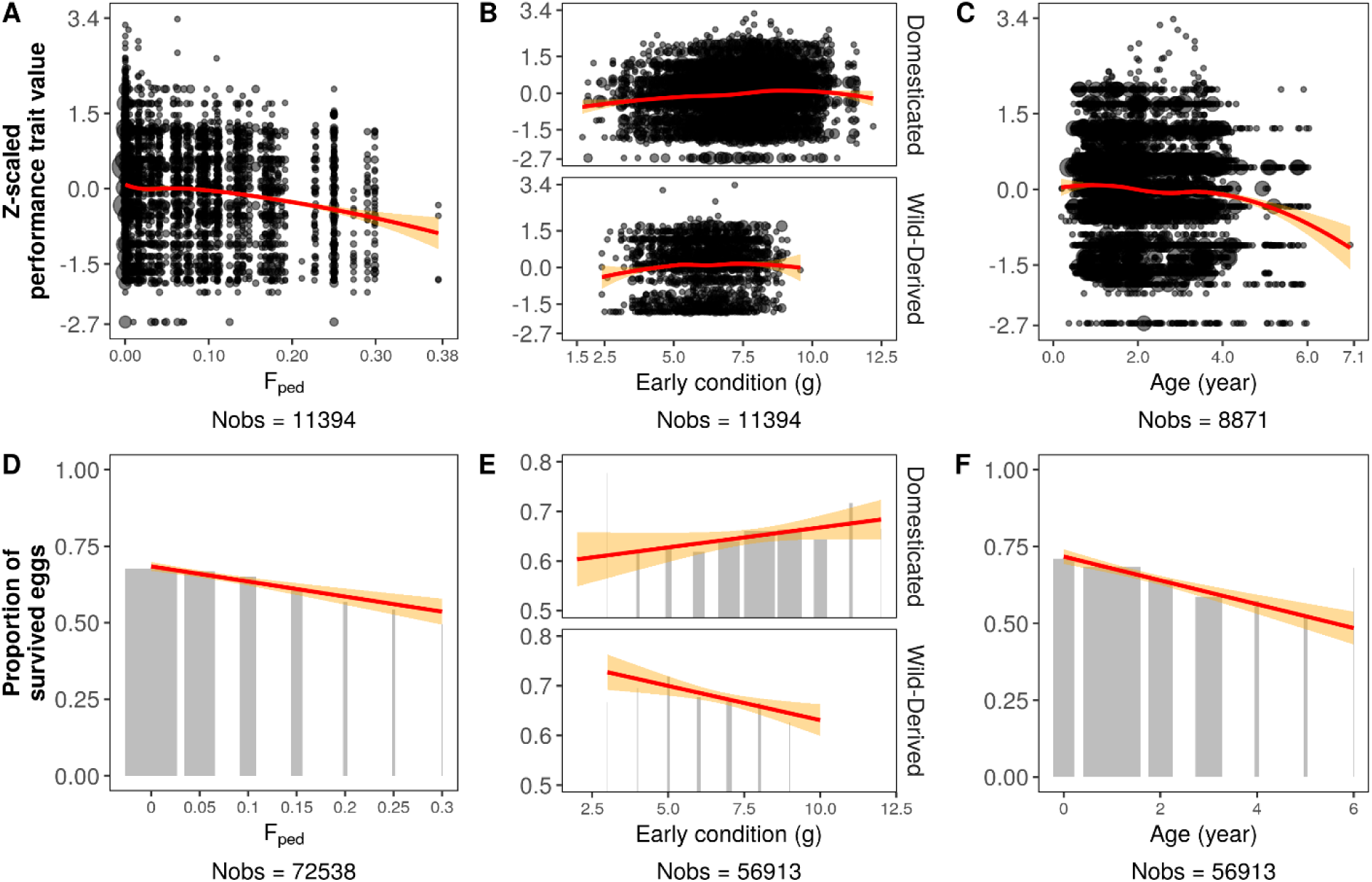
Reproductive performance traits (continuous or count traits in A-C, binomial traits in D-F) as a function of inbreeding coefficient (F_ped_; A, D), early condition (mass at day 8, separately for populations that differ in body size (B, E), and age (C, F). Clutch size, fecundity, siring success, seasonal recruits, and lifespan are continuous or count traits (Z-scaled), whereas the proportions of eggs fertilized, embryos survived, and nestlings survived are binomial traits. Note that these are composite figures of all effects that were examined (see fig. A2 for plots of single traits with absolute trait values), such that the fate of one embryo may be shown twice, once as a function of the embryo’s own F_ped_ and once as a function of its mother’s F_ped_ (hence the high sample sizes, N_obs_). The age category zero contains measurements until day 365. Red lines show smoothed regressions with 95% CIs, circle size (A-C) and bar width (D-F) reflect sample sizes.

**Figure 3.**
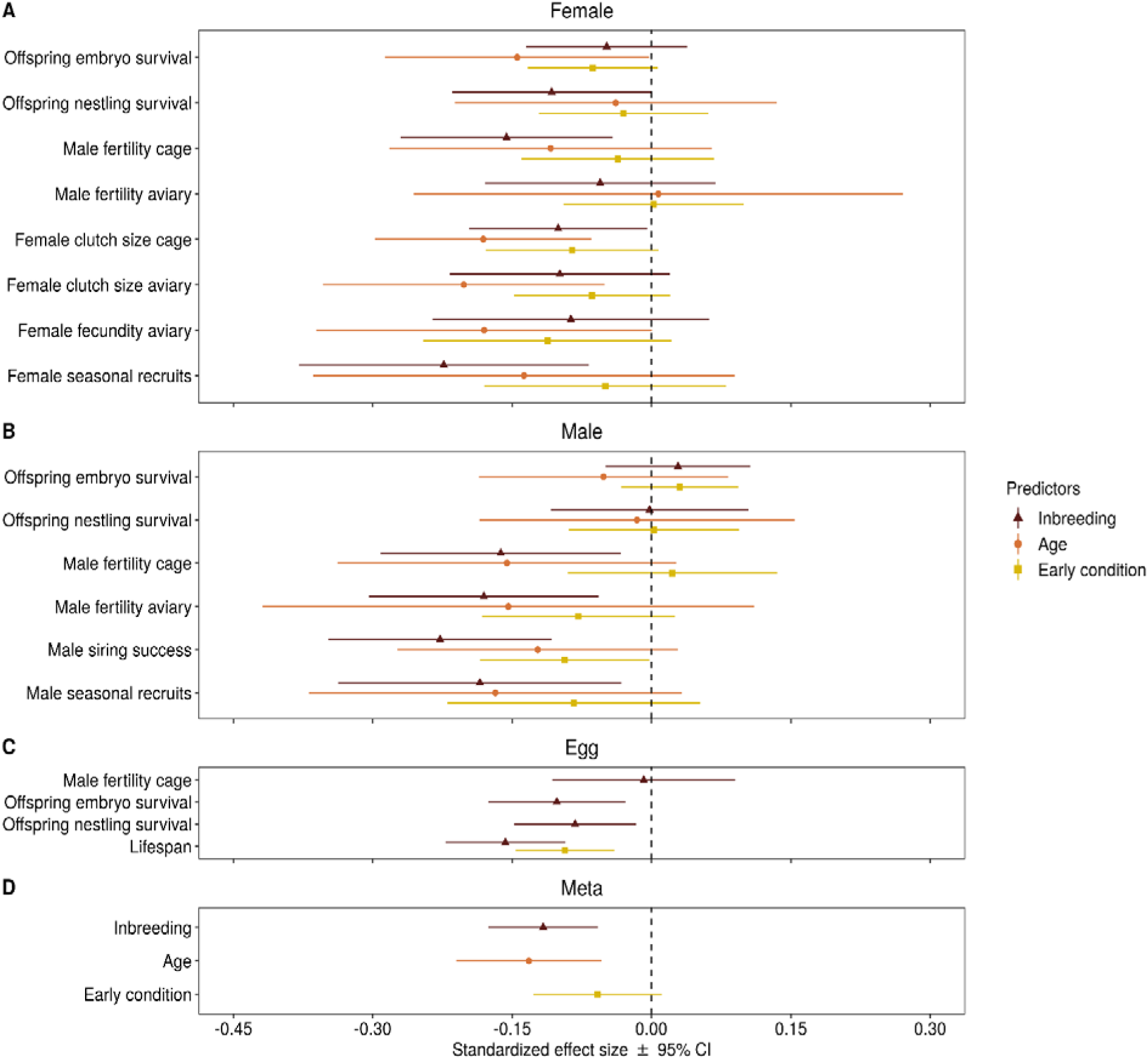
Standardized effect sizes with their 95% confidence intervals for inbreeding (F_ped_), age and early condition (mass at day 8) on zebra finch fitness components estimated in univariate Gaussian mixed-effect models where all response variables were measured at the level of individuals within seasons, and all measurements were Z-scaled (table A3). Note that the effect of inbreeding of the offspring on its own mortality was taken from egg-based models. Negative effects of condition indicate low fitness of relatively light-weight individuals at 8 days of age. Panels separate effects of condition, age and inbreeding of the female (A), the male (B), and the individual egg itself (C). Panel (D) shows the meta-summarized effect sizes for reproductive performance and lifespan (table A4). The X-axes indicate effect sizes in the form of Pearson correlation coefficients.

**Figure 4.**
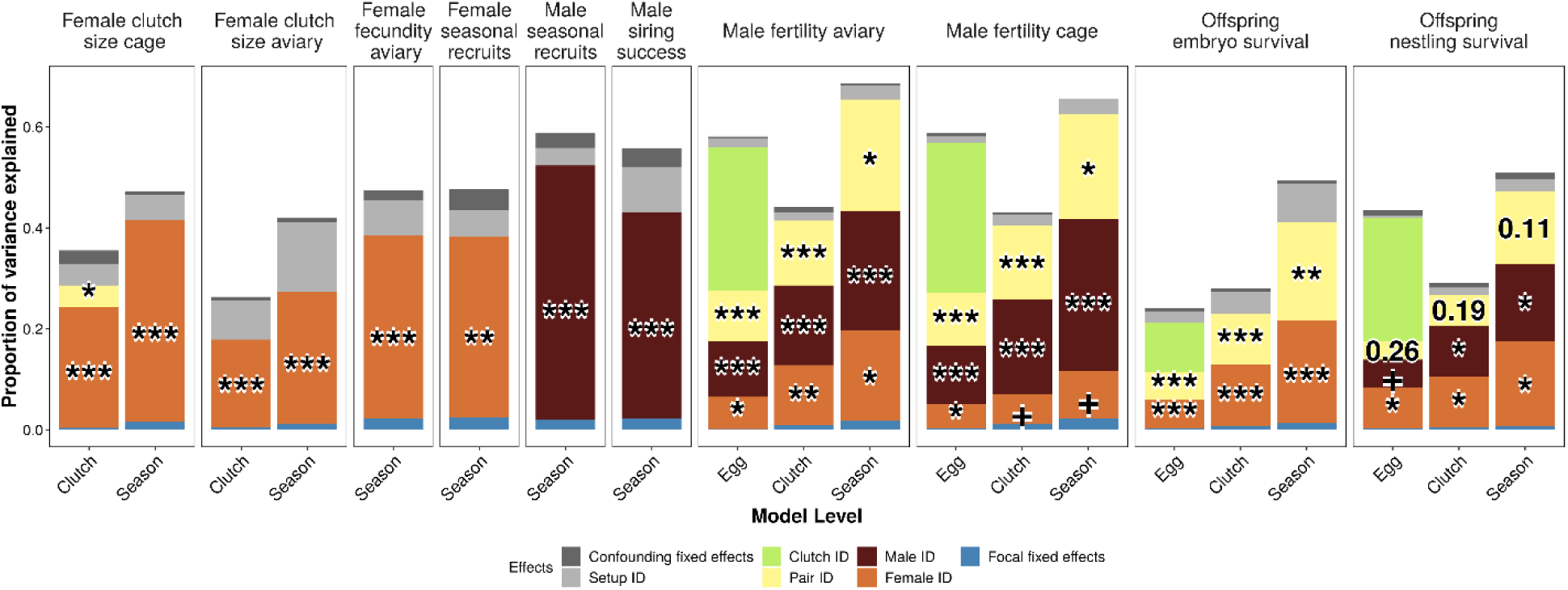
Variance components estimated in univariate Gaussian mixed-effect models (table A5). Each dependent trait is shown in a separate panel. Within panels, the x-axis separates models according to the unit of analysis, based on either egg fate (Egg), values per clutch (Clutch), or values per individual within a breeding season (Season). The y-axis indicates the proportion of variance explained by random effects after accounting for fixed effects. ‘Focal fixed effects’ refers to the total variance explained by inbreeding, age, and early condition combined. For the key variance components, numbers show non-significant P-values, otherwise ‘+’ indicates P < 0.1, ‘*’ indicates P < 0.05, ‘**’ indicates P < 0.001, and ‘***’ indicates P < 0.0001. Note that Models of ‘female clutch size aviary’, ‘female fecundity aviary’ and ‘female seasonal recruits’ were analyzed without ‘Male ID’ and ‘Pair ID’, and likewise ‘male seasonal recruits’ and ‘male siring success’ was analyzed without ‘Female ID’ and ‘Pair ID’ because not all birds form a pair bond; ‘Male ID’ explained no variance in models of ‘clutch size cage’ and ‘embryo survival’ while ‘Pair ID’ explained no variance in ‘clutch size cage’ model.

Meta-summarized effect sizes of inbreeding (r = -0.117 ± 0.024 SE) and age (r = -0.132± 0.032 SE) were similar in magnitude, and were about twice as large as the remarkably small effect of early condition (r = -0.058 ± 0.029 SE; fig. 3 and table A4). There was no significant difference between the categories male, female, and offspring in how strongly they were affected by these three factors (b ≤ 0.012 ± 0.028 SE, P = 0.63; table A4). Fitting trait (fitness component, 11 levels) as a random effect explained 1.5% of the variance in effect sizes (P = 0.02; table A4), suggesting that some components might be less sensitive than others (fig. 3; table A3). Female traits significantly predicted offspring survival and male fertility (independent of whether they were measured in a cage or in an aviary), whereas male traits showed no effect on offspring survival (fig. 3).

### Variance components and heritability

Variance components for all reproductive performance traits are shown in fig. 4 (see also table A4). Overall, individual reproductive performance traits were significantly repeatable (median R = 0.28, range: 0.15-0.50). Female reproductive performance traits (clutch size, fecundity, and female seasonal recruits) showed reasonably high repeatability for individual females (R ∼ 0.26- 0.40). Likewise, male fertility, male siring success, and male seasonal recruits were highly repeatable for individual males (R ∼ 0.24-0.50). Female reproductive traits from aviary-breeding were analyzed independently of whether the focal female had a partner or not (table 2), but female clutch size measured in a cage showed no contribution from the male partner or from pair identity. In contrast, male fertility depended on all three random effects, and was repeatable for males (R > 0.23, P < 0.0001), but less so for females (R <0.18, P < 0.1), and for the particular pair combinations (R < 0.23, P < 0.05). The model on embryo survival showed significant female and pair identity (genetic parents) effects that were similar in size (both R = 0.20, P < 0.0002), while genetic male identity explained no variance (fig. 4). In contrast, social female (R = 0.17, P = 0.017) and social male (R = 0.15, P = 0.039) identity explained significant amounts of the variance in nestling survival, while the effect of pair identity (parents that raised the brood) was less clear (R = 0.14, P = 0.11).

Reproductive performance traits and lifespan in general had low narrow-sense heritability (V_A_/V_ph_ ; Seewiesen: median h^2^ = 0.07; Bielefeld: median h^2^ = 0.11) and explained only a limited amount of the individual repeatability (V_A_/(V_A_+V_PE_); Seewiesen: median = 0.29; Bielefeld: median = 0.32; see all heritability estimates in tables A6-A7). Heritability estimates from the recently wild-derived population ‘Bielefeld’ were similar to those from the domesticated ‘Seewiesen’ population (for 9 traits measured in both populations; mean difference in h^2^ = 0.02, range: -0.10 - 0.13, meta-summarized difference after controlling for the uncertainty of each estimate: Δb < 0.0001; mean difference in V_A_/(V_A_+V_PE_) = 0.20, range: -0.13 - 0.68, meta-summarized difference: Δb = 0.0002; table A8).

### Additive genetic correlations

Reproductive performance traits were grouped into three classes: (1) aspects of male reproductive performance, (2) aspects of female reproductive performance and (3) aspects of offspring survival (table 2). Traits within each of these classes were on average positively correlated with each other at the additive genetic level (for the Seewiesen population, female traits: mean r_A_ = 0.66, P<0.0001; male traits: mean r_A_ = 0.67, P<0.0001; offspring survival traits: mean r_A_ = 0.36, P = 0.09; see fig. 5A). Results for the Bielefeld population are in fig. A6. The meta-summarized results are given in table A9 and all additive genetic correlation estimates are listed in tables A10-A11. Estimates of the additive genetic correlations from bivariate animal models using MCMCglmm are in fig. A7 (Seewiesen) and A8 (Bielefeld).

**Figure 5.**
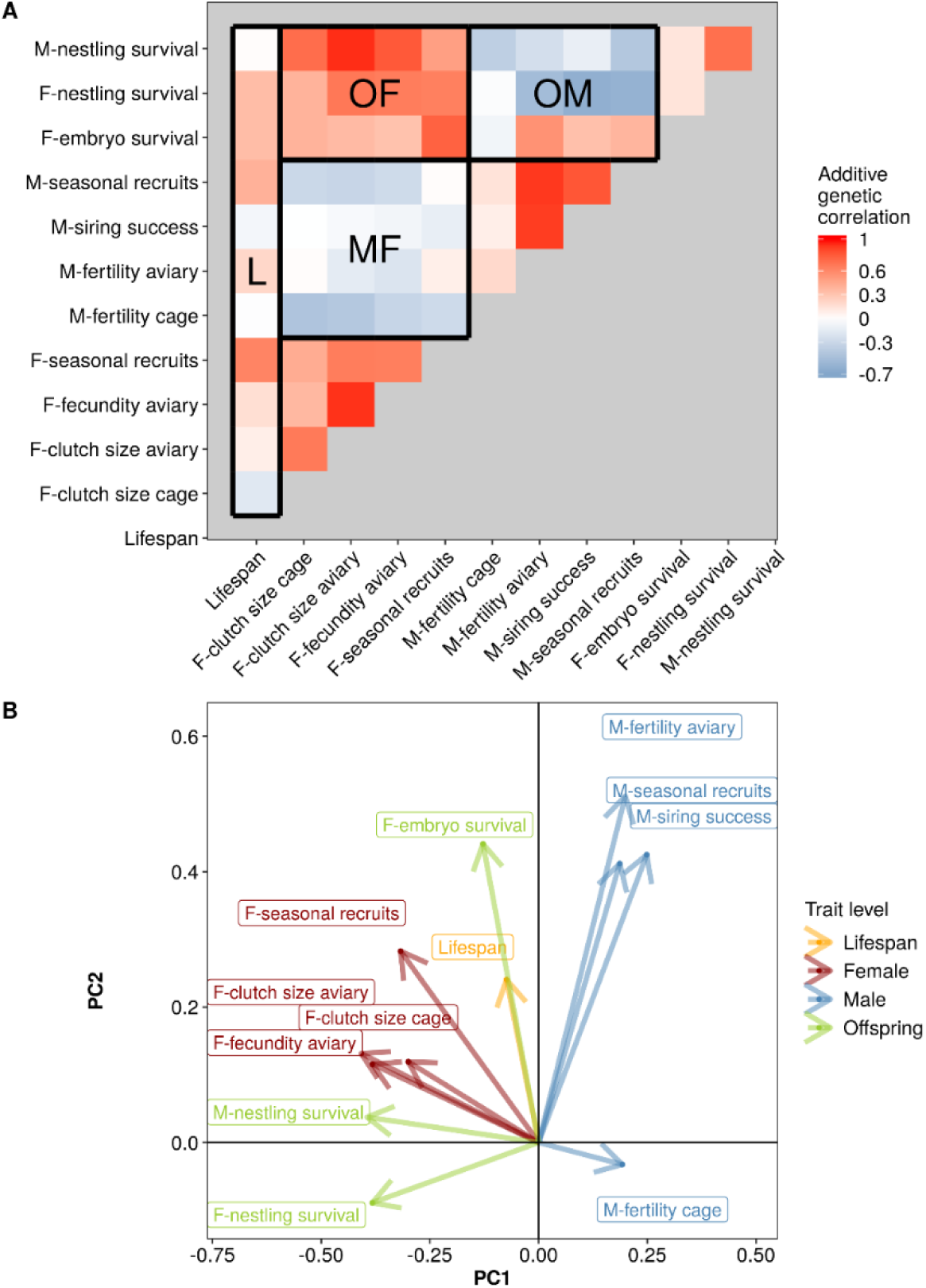
G-matrix of reproductive performance traits and lifespan estimated from multivariate animal models for the Seewiesen population (shown are estimates from VCE; for estimates of MCMCglmm bivariate models see fig. A7; see also figs A6 and A8 for estimates from the Bielefeld population; estimates are given in tables A10-A11). (A) Heatmap of additive genetic correlations between components of male (M), female (F), and offspring (O) fitness, and life span (L). Red indicates a positive genetic correlation between traits while blue indicates a negative correlation. Blocks marked in bold emphasize correlations between categories (e.g. MF stands for correlations between male and female fitness components). (B) The first two principal components of the G-matrix, showing eigenvectors of the 12 fitness components. Note that aspects of male fitness do not align with aspects of female and offspring fitness.

Male and female reproductive performance traits were weakly negatively correlated at the additive genetic level (mean r_A_ = -0.14, P = 0.04; see ‘MF’ in figs. 5A, A7A). Accordingly, the eigenvectors for male and female fitness traits were pointing into different directions (figs. 5B, A7B). This pattern was somewhat consistent between the Seewiesen and Bielefeld populations (see figs. A6, A8 for Bielefeld population). However, the negative correlation between male and female fitness traits was no longer significant when estimated by the bivariate animal models in ‘MCMCglmm’, and disappeared in the ‘Bielefeld’ dataset (table A9). The orientation of offspring survival traits relative to male and female fitness traits was less consistent. In the Seewiesen population, female fitness traits were positively correlated with offspring survival traits at the additive genetic level (mean r_A_ = 0.61, P < 0.0001), while male fitness traits were not aligned with offspring survival traits (mean r_A_ = -0.11, P = 0.24; fig. 5). In contrast, in the Bielefeld population, both female and male fitness traits were positively correlated with offspring survival traits (fig. A6). Lifespan tended to be positively correlated with all reproductive performance traits (Seewiesen: mean r_A_ = 0.19, P = 0.02; Bielefeld: mean r_A_ = 0.60, P = 0.0006; figs. 5, A6; table A9).

## Discussion

### Effects of inbreeding, age, and early condition

Many studies have shown that inbreeding depression significantly influences morphological, behavioral, and fitness-related traits in zebra finches (Bolund et al. 2010a; Forstmeier et al. 2012; Hemmings et al. 2012; Opatová et al. 2016), and in other species (Amos et al. 2001; Reed and Frankham 2003; Williams et al. 2003; Michaelides et al. 2016). This study confirms that inbreeding negatively influenced all phases of offspring survival, reproductive performance and lifespan. We found that the level of inbreeding of both genetic parents negatively influenced egg fertility, suggesting that this is not only a matter of sperm functionality (Opatova et al. 2016), but also of female reproductive performance (e.g. egg quality or copulation behavior). Male and female fitness estimates (seasonal recruits) were most strongly affected by inbreeding (fig. 3), presumably because the successful rearing of offspring to independence requires proper functionality at every step of reproduction.

Age effects on reproductive performance typically show an initial increase in performance both in short- and in long-lived species (e.g. in great tits Parus major (Bouwhuis et al. 2009), Langur monkeys Presbytk entellus (Harely 1990), red deer Cervus elaphus (Pemberton et al. 2009), albatrosses Diomedea exulans (Lecomte et al. 2010), and bustards Chlamydotis undulata (Preston et al. 2011)). Interestingly, in our captive zebra finches we found that reproductive performance (especially male fertility, female clutch size, fecundity, and female effects on embryo survival) did not show an initial increase after birds reached sexual maturity at about 100 days of age (figs. 2C, F, A2). This could be because zebra finches are short-lived opportunistic breeders that reach sexual maturity earlier compared to most other birds (Zann 1996). Thus, zebra finches might have been selected to perform best early on. Alternatively, this effect may not be present in the wild, where experience might play a more important role in determining reproductive success.

Over the past decades, numerous studies focused on how early developmental conditions affect behavior, life history, and reproductive performance later in life (Tschirren et al. 2009; Rickard et al. 2010; Boersma et al. 2014). Here, we show that even dramatic differences in early growth conditions of surviving offspring (see range of x-axis in fig. 2B), have remarkably small (though statistically significant) effects on adult reproductive performance.

Overall, the proportion of variance explained by inbreeding, age, and early condition (characteristics of conditions) was less than 3% (fig. 4; table A4). This indicates that individuals’ robustness against poor conditions appears more noteworthy than their sensitivity. As will be discussed in the following paragraphs, the majority of the individual repeatability in reproductive performance cannot be explained by such individual characteristics.

### Repeatability and heritability of reproductive performance

Individual zebra finches were remarkably repeatable in their reproductive performance. Our variance-partitioning analysis showed that infertility is largely a male-specific trait, whereas embryo and offspring survival are mostly related to female identity (fig. 4; table A4). The effects of pair identity on infertility and offspring mortality may reflect behavioral incompatibility, while the pair effect on embryo mortality more likely reflects genetic incompatibility (Ihle et al. 2015).

Although male and female zebra finches are highly repeatable in their reproductive performance, the heritability of fitness traits was low and similar between the recently wild-derived Bielefeld population and the domesticated Seewiesen population. Overall, our findings indicate that there are some additive genetic components underlying zebra finch reproductive performance.

### Evidence for sexually antagonistic pleiotropy

Some of the standing additive genetic variance in reproductive performance could be maintained by intra-locus sexual antagonism between male fitness traits and female (and offspring) fitness traits. This has for example also been suggested in studies on Drosophila (Innocenti and Morrow 2010) and on red deer (Foerster et al. 2007). We found that male fertility, siring success and seasonal recruitment were overall negatively correlated with female fitness and offspring survival traits, suggesting that alleles that increase male fitness tend to reduce female and offspring fitness (fig. 5). In contrast, lifespan and reproductive performance tended to be positively correlated at the additive genetic level, which is suggestive of some overall ‘good gene variation’ in our population (fig. 5). Some words of caution should be added to these observations. VCE6 (figs. 5, A6) yielded higher absolute values of estimates than those calculated with the R functions ‘PedigreeMM’ (heritability estimates only) and ‘MCMCglmm’ (see appendix figs. A7-A8; also see tables A6-A7, A10-A11). Nevertheless, the additive genetic correlation estimates are highly correlated between the two methods (r > 0.7, P < 0.0001; see tables A10-A11). Estimating genetic correlations between traits with low heritability requires large datasets, especially on additive genetic correlations of between-sex reproductive performance where the traits of male fertility and female clutch size in cages are missing (N performance traits: Seewiesen = 12, Bielefeld = 9; N birds have at least one entry of reproductive performance data: Seewiesen = 2346, Bielefeld = 1134; hence these results are presented in the online appendix only, fig. A6). Despite this lack of power in our second largest data set of population Bielefeld, its overall orientation of traits in the additive genetic variation space of the principle components PC1 and PC2 is very similar to population Seewiesen (note that lifespan is in the center of all fitness traits and that aspects of female fitness do not align with male fitness in figs. 5*B*, *A6B, A7B, A8B*).

Individual repeatability of fitness-related traits could arise from permanent environmental effects (e.g. early developmental conditions and long-lasting diseases) or from genetic effects. However, while food-shortage experienced during early development (reflected in body mass at 8 days old) strongly predicted nestling mortality (our unpublished data), it only explained <1% of variation in reproductive performance (mean r = -0.058, figs. *2B, E*, 3 and also 4). Additionally, our captive zebra finches were raised and kept in a controlled environment with no obvious diseases detected. Additive genetic effects explained only about 30% of the large remaining unexplained individual repeatability in fitness-related traits, suggesting that reproductive performance might be (predominantly) dependent on genetic effects of local over- or under- dominance and epistasis, i.e. incompatibility between loci. For instance, high levels of reproductive failure could be maintained when alleles show non-additive effects, with selection favouring the heterozygous genotype (see e.g. Sims et al. 1984; Grossen et al. 2012). In the zebra finch, males that are heterozygous for an inversion on the Z chromosome produced fast-swimming sperm and sired more offspring (Kim et al. 2017; Knief et al. 2017). Epistatic effects that involve several genes (e.g. incompatibility between nuclear loci, or between mitochondrial and nuclear genomes) could be evolutionary stable when certain combinations of genotypes perform better than others, especially when combined with overdominance (Avent and Reid 2000; Arntzen et al. 2009; Hermansen et al. 2014; Knegt et al. 2016; Baris et al. 2017; Stryjewski and Sorenson 2017).

Infertility, as one of the main and puzzling sources of reproductive failure, behaved as a male-specific trait that may in part also depend on behavioral compatibility between pair members (reflected in copulation behavior) and in part on the male’s genotype at sexually antagonistic loci. The intrinsic male fertility, measured in a cage, i.e. in the absence of sperm competition, correlated negatively with all female and offspring survival traits at the additive genetic level (‘sexual antagonism’; median r_A_ = -0.30, range: -0.45 to -0.01; fig. 5; table A10). In contrast, in the presence of sperm competition (aviary breeding), high male fertility, siring success and seasonal recruitment should also be influenced by the competitive ability of the individual and this could explain why these traits correlated positively with lifespan and trade-off less with female traits and offspring rearing ability at the additive genetic level (figs. 4, A6; tables A10-A11).

Embryo mortality, another main source of reproductive wastage, mostly depended on the identity of the genetic mother and the identity of the genetic pair members. A previous study using cross-fostering of freshly laid eggs also showed that embryo mortality is a matter of the genetic parents rather than the foster environment (Ihle et al. 2015). The female component suggested an overall female genetic quality effect, yet with limited heritability (pointing towards dominance variance or epistasis). The effect of the combination of parents on embryo mortality might reflect an effect of the genotype of the embryo itself, possibly involving multi-locus incompatibilities.

## Conclusions

Our results suggest that sexually antagonistic pleiotropy between male and female fitness plus offspring rearing traits may maintain some of the existing additive genetic variation in reproductive performance traits in captive zebra finches. Additionally, there appears to be some ‘good gene’ (heritable) variation among reproductive performance traits and individual lifespan, which suggests an ongoing adaptation to the captive environment. We found that the level of inbreeding, age and – to a lesser extent – early rearing conditions predicted a small, but statistically significant amount of variation in individual reproductive performance and lifespan. However, those three effects were so small that they cannot be the main causes of reproductive failure. Although individual zebra finches were moderately repeatable in their reproductive performance, the heritability of those traits was low. Overall, our results suggest that alleles that have additive effects on fitness might be maintained through sexually antagonistic pleiotropy, and the major genetic causes of reproductive failure might be determined by genetic incompatibilities or local dominance effects.

## Acknowledgements

We thank M. Schneider for molecular work, H. Schielzeth, E. Bolund, M. Ihle, U. Knief, C. Burger, S. Janker, J. Schreiber, T. Aronson, and A. Edme for help with data collection, T. Birkhead, T. Krause, J. Lesku, and K. Roberts for providing birds, and S. Bauer, E. Bodendorfer, J. Didsbury, A. Grötsch, J. Hacker, A. Kortner, J. Minshull, P. Neubauer, C. Scheicher, F. Weigel, and B. Wörle for animal care and help with breeding.

